# Structural Basis of RNA Cap Modification by SARS-CoV-2 Coronavirus

**DOI:** 10.1101/2020.04.26.061705

**Authors:** Thiruselvam Viswanathan, Shailee Arya, Siu-Hong Chan, Shan Qi, Nan Dai, Robert A. Hromas, Jun-Gyu Park, Fatai Oladunni, Luis Martinez-Sobrido, Yogesh K. Gupta

## Abstract

The novel severe acute respiratory syndrome coronoavirus-2 (SARS-CoV-2), the causative agent of COVID-19 illness, has caused over 2 million infections worldwide in four months. In SARS coronaviruses, the non-structural protein 16 (nsp16) methylates the 5’-end of virally encoded mRNAs to mimic cellular mRNAs, thus protecting the virus from host innate immune restriction. We report here the high-resolution structure of a ternary complex of full-length nsp16 and nsp10 of SARS-CoV-2 in the presence of cognate RNA substrate and a methyl donor, S-adenosyl methionine. The nsp16/nsp10 heterodimer was captured in the act of 2’-O methylation of the ribose sugar of the first nucleotide of SARS-CoV-2 mRNA. We reveal large conformational changes associated with substrate binding as the enzyme transitions from a binary to a ternary state. This structure provides new mechanistic insights into the 2’-O methylation of the viral mRNA cap. We also discovered a distantly located ligand-binding site unique to SARS-CoV-2 that may serve as an alternative target site for antiviral development.

## Introduction

The massive global pandemic with high morbidity and mortality makes SARS-CoV-2 one of the deadliest virus in recent history^*1*^. To develop effective therapies, we need a better understanding of the mechanisms that permit the virus to invade cells and evade host immune restriction. SARS-CoV-2 is an enveloped, positive-sense single-stranded β-coronavirus with a large, complex RNA genome^*2*^. To hijack the host translation machinery for propagation, enzymes encoded by the genome of coronaviruses (CoVs) modify the 5’-end of virally encoded mRNAs by creating a cap^*3*^. RNA capping in CoVs involves activities of several nonstructural proteins (nsps): nsp13, a bifunctional RNA/NTP triphosphatase (TPase) and helicase; nsp14, a bifunctional 3’→5’ mismatch exonuclease and mRNA cap guanine-N7 methyltransferase; nsp16, a ribose 2’-O methyltransferase; and an elusive guanylyl transferase^*4–7*^.

Nsp16 forms an obligatory complex with nsp10 to efficiently convert client mRNA species from the Cap-0 (^me7^G_o_ppp**A**_**1**_) to the Cap-1 form (^me7^G_o_ppp**A**_**1m**_), by methylating the ribose 2’-O of the first nucleotide (usually adenosine in CoVs) of the nascent mRNA using SAM (S-adenosyl methionine) as the methyl donor ^*4, 8*^. This Cap-1 form serves to avoid induction of the innate immune response^*9–11*^. Hence, ablation of nsp16 activity should trigger an immune response to CoV infection and limit pathogenesis^*9, 10*^. It has been shown that live vaccination with nsp16-defective SARS-CoV-1 or an immunogenic disruption of the nsp16-nsp10 interface protects mice from an otherwise lethal challenge^*12, 13*^, making nsp16/nsp10 an attractive drug target.

Crystal structures of SARS-CoV-1 nsp16/nsp10 in complex with SAM/SAH or sinefungin, but without an RNA substrate, have been reported ^*4, 8, 14*^, so that key information about the catalytic mechanism of mRNA capping in CoVs, and SARS-CoV-2 in particular, is still missing. To understand the determinants of RNA cap modification and help guide the development of SARS-CoV-2 antiviral therapies, we have now succeeded in solving the high-resolution structure (to 1.8 Å resolution) of SARS-CoV-2 nsp16/nsp10 in complex with the methyl donor (SAM) and its target, the RNA cap analog ^me7^G_o_ppp**A**_**1**_. Moreover, we co-crystallized this complex with a series of FDA-approved drugs and found that one such compound, adenosine, associates with a probable allosteric ligand binding site not present in the nsp16/nsp10 orthologues from other SARS or pan-coronavirus strains. Thus, we provide a plausible novel framework from which SARS-CoV-2 inhibitors may be developed.

## Results

We co-purified full-length SARS CoV-2 nsp16 and nsp10 proteins as a complex from *E. coli*, mixed with nucleoside drugs such as adenosine or 5’-methylthioadenosine and subjected to crystallization screenings (see Methods for details). We anticipated that these drugs may occupy the binding site of SAM due to common features of both, a purine ring plus ribose. Since nucleoside analogues exhibit antiviral activity^*15–17*^, a nsp16/nsp10/adenosine complex could then serve as a starting point for medicinal chemistry and perhaps rapid translation. Next, we soaked these crystals with a cap analogue ^me7^G_o_ppp**A**_**1**_ representing the Cap-0 state of the RNA cap, the substrate of this complex. We determined the ternary structure of nsp16/nsp10 by molecular replacement using the SARS-CoV-1/SAM binary complex (PDB ID 3R24)^*8*^ structure as a template (Supplementary Table 1). The initial difference maps indicated electron densities of SAM and ^me7^G_o_ppp**A**_**1**_ cap in their respective binding pockets. Even though SAM was not included in co-crystallization or soaking experiments, its electron density was unambiguously identified in difference maps (Fig. 1, 2, Supplementary Fig. 2A). This observation is consistent with the co-purification tendency of SAM during isolation from expression hosts, as seen previously with NS5 of ZIKA and VP39 of vaccinia virus cap 2’-O MTases ^*18, 19*^. Unexpectedly, we observed electron density on the opposite face of nsp16, >25 Å away from the catalytic pocket that could fit adenosine (present in our crystallization mixture) reasonably well (Figure 1, Supplementary Fig. 2, 3). Further examination showed that this adenosine-binding pocket was partially composed of amino acid residues unique to SARS-CoV-2 (Supplementary Fig. 1)

**Fig. 1.**
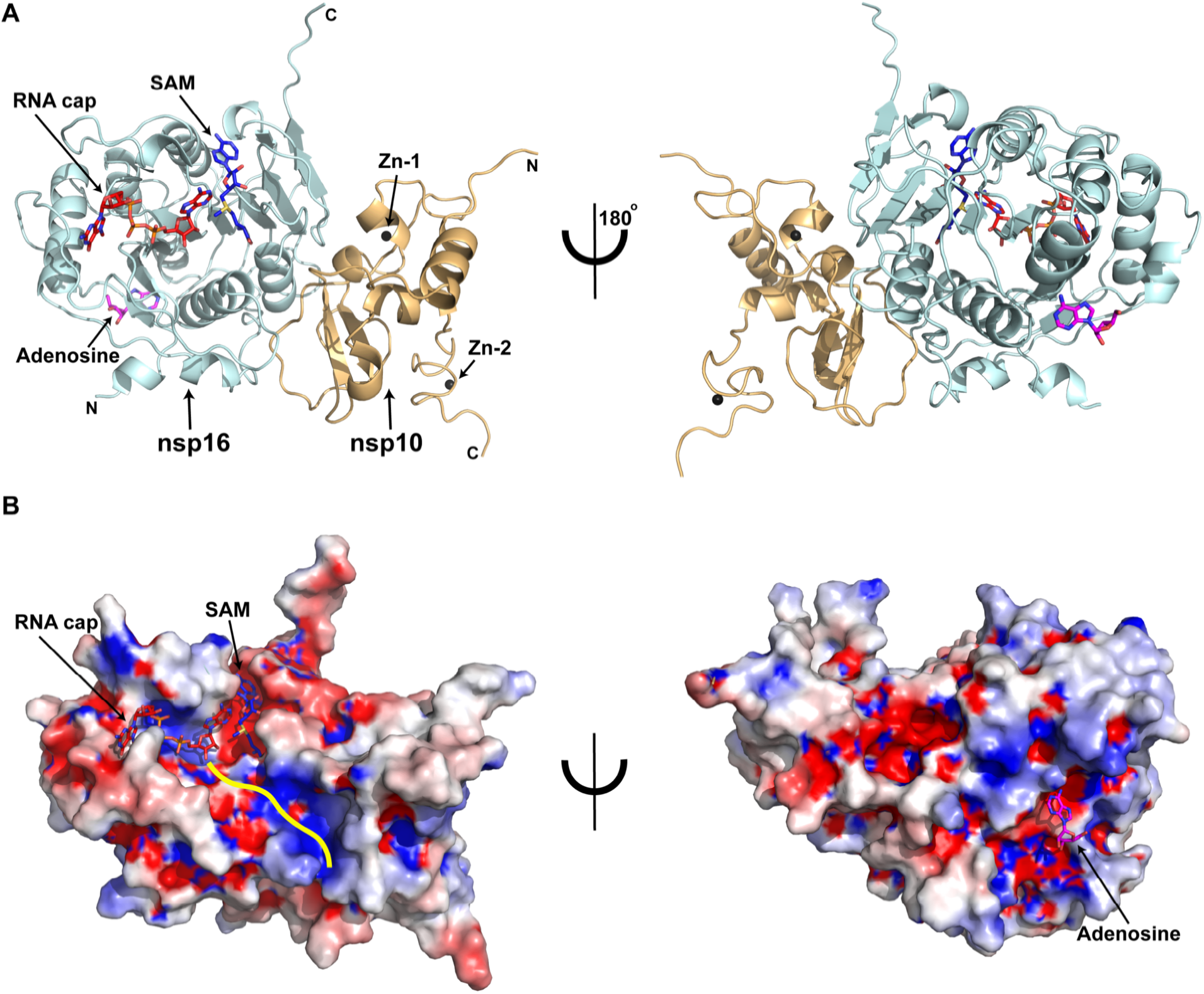
Overall structure of the SARS-CoV-2 nsp16/nsp10 ternary complex. A) Subunit arrangement of nsp16 (cyan) and nsp10 (orange cartoons) proteins with respect to RNA cap (red), and SAM (blue). Proteins, nucleotides are shown in cartoon and stick modes, respectively. Black spheres, Zn^2+^ atoms bound to nsp10; Magenta stick, adenosine bound to nsp16. B) Electrostatic surface representation of nsp16/nsp10 with saturated blue and red at +5 kT/e and −5 kT/e, respectively. A yellow line represents a tentative path for downstream RNA sequence calculated by superposing the target bases in current, VP39, and Dengue NS5 ternary complexes.

**Fig. 2.**
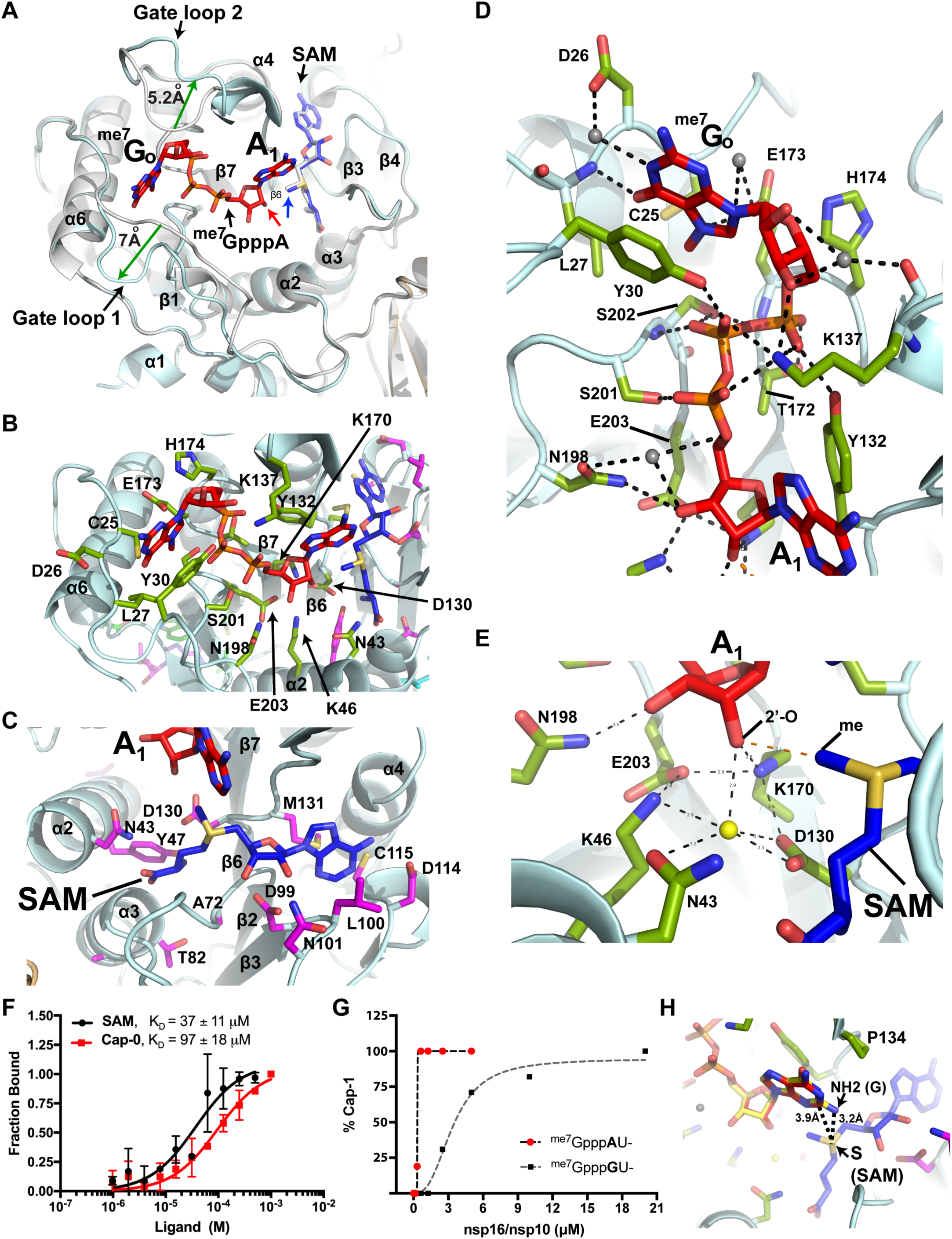
Binding modes of RNA cap analogue and SAM, and mechanism of methyl transfer. **A)** Overlay of a binary (SAM-bound; grey cartoons) and ternary (SAM, RNA cap-bound; light cyan) complexes shows outward motions (green arrows: 7 Å in gate loop1, and 5.2 Å in gate loop 2) after RNA cap binding to nsp16. RNA cap, red; SAM in ternary complex, blue; SAM in binary complex, grey stick. **B)** green; nsp16 residues that interact with RNA cap. **C)** magenta; nsp16 residues that contact with SAM. **D)** A close-up view of Cap-nsp16 interactions reveals a network of hydrogen bonding with successive phosphates, ^me7^G_o_ and A_1_ nucleotides of cap. Water, grey spheres, h-bonds, dashed lines. **E)** A water (yellow sphere) coordinates with the target 2’-O atom of A_1_, and catalytic triad residues and N43. The methyl group of SAM is positioned for direct *in line* attack from the 2’-O. **F)** Binding isotherms and fitting of data for nsp16 binding to RNA cap (^me7^GpppA) and SAM. **G)** The 2’-O methyltransferase activity measured as percentage of Cap-0 to Cap-1 conversion is plotted against nsp16/nsp10 protein concentration. Higher enzymatic activity is observed on an RNA substrate with **A** (red circles) as the target base for 2’-O methylation (N_1_), compared to an identical RNA but with **G** (black square) as N_1_ or initiating nucleotide. **H)** Guanine base (yellow stick) is modeled at N_1_ position of cognate adenine (red stick). The N2 amine of guanine intrudes into the SAM pocket and may be repelled by positively charged sulfur of SAM (blue stick).

**Fig. 3.**
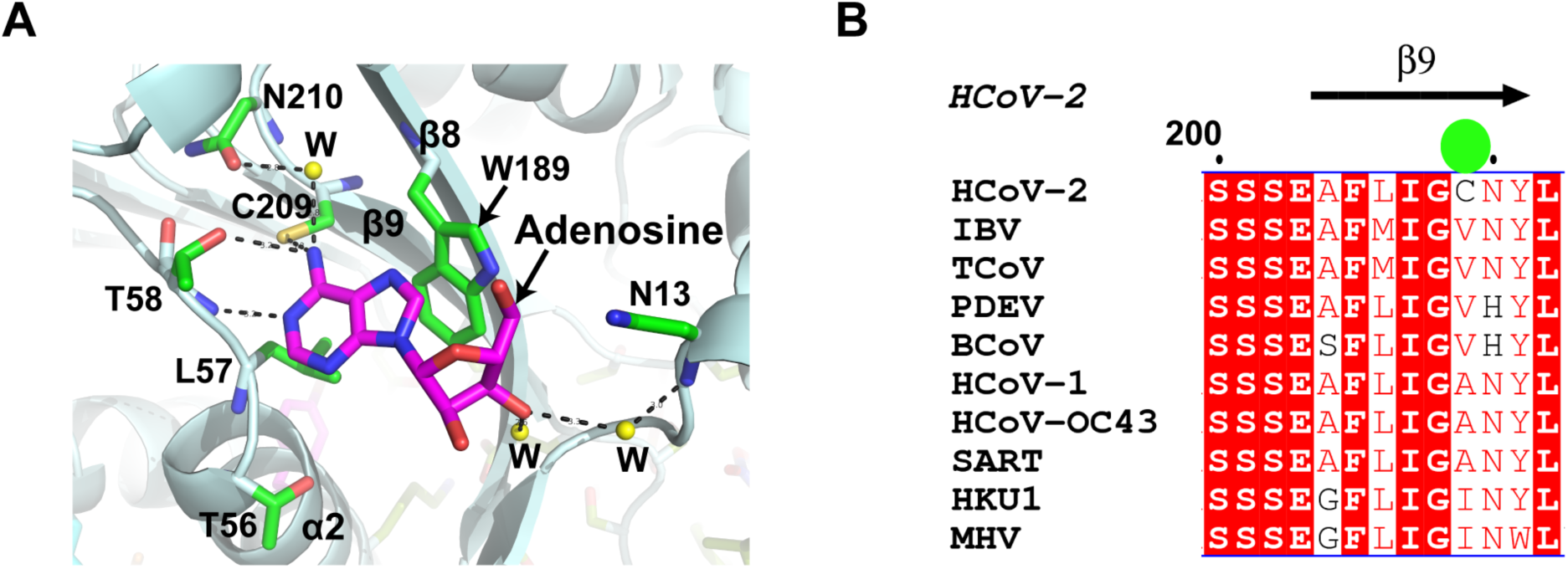
Architecture of a putative allosteric site in nsp16. A) nsp16 residues that participate in adenosine (magenta) binding are shown as green sticks; water molecules are shown as yellow spheres. B) Cys209 (green sphere) in the β9 strand, which is present in SARS-CoV-2, but no other CoV strain.

### Overall structure

Nsp16 adopts a canonical S-adenosyl methionine (SAM)-dependent methyltransferase (SAM-MTase) fold^*20*^ with slight variations ^*8, 14*^. Its protein sequence displays a sequential order of secondary structural element: •β1α1α2β2α3β3β4•β5β6•α4β7α5β8•β9α6•β10•α7β11β12, wherein the ‘•’ denotes a 3_10_-helix (Fig. 1, Supplementary Fig. 1). The nsp16 MTase fold consists of a centrally located twisted β sheet of eight strands flanked by two alpha-helices on one side and three helices on the other. The β sheet displays a continuum of four antiparallel (β1β8β9β7) and four parallel (β6β2β3β4) β strands. The cap analogue substrate is situated within the confluence of these two halves. Loops emanating from strands β9, β7, and β6 form a deep groove in the center to accommodate the RNA cap, whereas the methyl donor SAM is bound within a cavity formed by the loops originating from strands β6 and β2. The protein chain emerging from helix α4 runs across this groove and folds into a subdomain (•β10•α7β11β12) that stabilizes the bottom portion of nsp16. The adenosine binding pocket is at the back of the catalytic pocket, ~ 25 Å away (Fig. 1).

Nsp10 is a 139 amino acid long zinc-binding protein that stimulates the enzymatic activity of nsp16 ^*4, 8, 14*^. We traced all functional regions known for protein-protein and protein-metal binding in nsp10, with the exception of the N-terminal 17 residues that appear to be disordered in our structure, but form an α-helix in the binary state in the absence of a cap structure ^*4, 8, 14*^. Nsp10 adopts a structural fold with two distinct Zn-binding modules, including a gag-knuckle-like fold ^*21*^. Binding of the RNA cap did not induce any major conformational change in nsp10.

### RNA substrate binding

We compared our ternary structure with bound substrate to that of SARS-CoV-1 nsp16/nsp10 bound to SAM (PDB ID 3R24), but without substrate. While the cores of the nsp16 and nsp10 proteins remain largely unperturbed (with an RMSD of 1.11 Å for 292 Cα atoms between the prior CoV-1 structure and our own), we found significant deviations in two regions of nsp16 that constitute the substrate binding pocket. We refer to these regions as gate loop 1 (amino acids 20-40) and gate loop 2 (amino acids 133 – 143) (Fig. 2). The binding of the Cap-0 substrate results in an ~180° outward rotation of gate loops 1 and 2 by 7 Å and 5.2 Å, respectively compared to their positions in the binary structure. The widening of the pocket that results allows accommodation of the RNA cap substrate, and engages the *N*^*7*^-methyl guanosine base of Cap-0 (^me7^G_o_) in a deep groove formed by residues of gate loop 1 on one side and gate loop 2 on the other (Fig. 2A, B, D, F). The loop region immediately downstream of Pro134 that is part of gate loop 2 also flips ~180º in the ternary complex to accommodate the A_1_ nucleotide (Fig. 2A). Gln28 and Asn29 (gate loop 1) orient outward, whereas the side chain of invariant Tyr30 rotates inward, thereby forming a cleft to stabilize the purine ring of ^me7^G_o_ through a π-π stacking interaction.

The opposite face of the ^me7^G_o_ purine ring is stabilized by a H-bond between terminal oxygen of Glu173 and the positively charged N7 of ^me7^G_o_. The methyl group of ^me7^G_o_ is stabilized by hydrophobic interactions with Cys25 (gate loop 1) and Ser202 from the loop connecting the β8 and β9 strands. These specific interactions with the positively charged N7 and the methyl moiety of the ^me7^G_o_ base would confer substrate selectivity against the uncharged G_o_pppA ^*4*^. In gate loop 2, Asn138 rotates outward and the side chain of invariant Lys137 rotates ~180º inward and forms stabilizing electrostatic interactions with all 3 phosphates of the Cap-0 structure and the 3’-O of the ^me7^G_o_ ribose. Tyr30 and Lys137 form a partial enclosure to restrict the movement of the terminal residue (^me7^G_o_) of the RNA cap (Fig. 2B, D). The backbone of gate loop 1 runs perpendicular to the purine ring of ^me7^G_o_ such that the backbone amide of Leu27 forms a hydrogen bond with O6 of ^me7^G_o_ to confer specificity for guanine. The ribose sugar of ^me7^G_o_ is sandwiched by the sidechains of Lys137 and His174 in gate loop 2 (Fig. 2B, D). The three phosphates of ^me7^G_o_ppp**A**_**1**_ are stabilized through H-bonds with residues located in the loop regions emanating from the β6 (Tyr132, Lys137), β7 (Thr172 and His174) and β8 (Ser 201 and Ser 202) strands (Fig. 2D). By comparing our structure with those for the 2’O-MTases of Dengue and Vaccinia viruses, we note that the distance between N_1_ and ^me7^G_o_ bases remains unchanged (~10 Å), and the SAM/SAH and N_1_ in all 3 structures overlay well, but the terminal ^me7^G_o_ base of CoV-2 RNA cap rotates ~180º (compared to Dengue virus) around the γ-phosphate to assume a radically different orientation, suggesting a diverse geometric arrangement of the RNA cap in RNA/DNA viruses (Supplementary Fig. 2B).

### Methyl donor (SAM) binding

SAM occupies the same pocket and orients in a similar manner as in the binary complex (PDB ID 3R24). Several loops emanating from the C-terminal ends of the four parallel β strands (β2, β3, β4, and β6) of the central β sheet form a deep groove to accommodate SAM through an extensive network of electrostatic, hydrophobic, and van der Waals interactions (Fig. 2C, F). In contrast to the substrate pocket, which is largely basic, the SAM binding pocket in nsp16 is enriched by negatively charged residues (Fig. 1B). The side chains of Asp99 and Asn101 from strand β3 form hydrogen bonds with terminal oxygens of the ribose sugar, whereas the purine ring of SAM is partially circled by side chains of Leu100 (β3) Asp114, Cys115 (β4), and Met131 (β6), and Trp149. Asp114 makes a base-specific interaction with SAM via a hydrogen bond with the N6 amino group of the purine ring. A four amino acid long stretch (Met131 to Pro134) that corresponds to gate loop 2 intrudes into the SAM and cap binding pockets and separates the two purine rings of SAM and A_1_ (the target adenine) by ~10 Å. This separation favorably orients the ribose sugar of A_1_ and the donor methyl group of SAM for 2’-O methylation. The carboxy end of SAM is stabilized by charged interactions from the side chains of Asn43 and Tyr47 and the backbone amide of Gly81 (Fig. 2C).

### Methylation of target adenine of RNA cap

Based on the crystal structures of SAM and SAH-bound SARS-CoV-1 nsp16/nsp10 complexes, a catalytic mechanism that involves a tetrad consisting of the invariant Lys46, Asp130, Lys170, and Glu203 has been proposed ^*8, 14*^. Our ternary complex has allowed us to examine the model in the presence of the Cap-0 substrate. The catalytic pocket adopts a conical shape where residues of the catalytic tetrad circle a water molecule at the ‘bottom’ of the cone, whereas the target atom (2’-O) of the A_1_ nucleotide resides at the ‘tip’ of the cone. In addition, two invariant asparagine residues - Asn43 and Asn198 - outside of the catalytic tetrad stabilize the water molecule and the 3’-end of the target nucleotide (A_1_) in the RNA cap, respectively. The methyl group of SAM is positioned 3 Å from the 2’-OH of A_1_, available for a direct *in line* attack from the 2’-OH (Fig. 2C, E). Supporting this concept, nsp16/nsp10 showed robust 2’-O methyltransferase activity. We also observed marked reduction in activity when the initiating nucleotide (N_1_) was changed from the cognate adenine to non-cognate base guanine (Fig. 2G). Similar observations were made by previous biochemical studies^*8*^, but the structural basis of this base discrimination was not clear. There is no base specific contact exists for the N_1_ base in ternary structure. However, modeling of a guanine (G) base at this position provides some clues. The Pro134 resides at Van der Waals distance from purine ring of the N_1_ base. A guanine at N_1_ will not alter this interaction but the N2 amino group of guanine intrudes into the SAM pocket thereby reducing the distance between guanine and SAM by ~0.7 Å. As a result, positively charged sulfur of SAM may repel purine ring of guanine and thus re-orients its target sugar 2’-OH in an unfavorable position for methyl transfer (Fig. 2H).

An earlier study posits that the target 2’-OH of the ribose in a ternary complex would initially occupy the position of the water, allowing Asp203 and Lys170 to lower the pKa of the ε-amino group of Lys46. As a result, Lys46 would become a deprotonated general base and activate the 2’-OH to attack the methyl group of SAM^*14*^ with Asp130 stabilizing the positive charge of the methyl group^*14*^. In our ternary structure, this water molecule resides 3 Å from the target 2’-OH of the A_1_ base ribose and remains unperturbed in the ternary structure of SARS-CoV-2 (present study) and binary structures (SAM/SAH-bound) of SARS-CoV-1^*8, 14*^ nsp16. If the ribose were to occupy the position of water as suggested ^*14*^, the adenine ring of A_1_ would sterically clash with SAM. In addition, Lys170 is closer to the 2’OH than Lys46 (Fig. 2E) and is held in place by charged interactions from the side chains of Asp130 and Glu203 from both sides. Consistent with this logic, Lys170 in our ternary complex structure spatially matches with that activating the 2’-OH in other viral 2’-O MTases (e.g., Lys180 of dengue virus NS5^*22*^, and Lys175 of vaccinia virus VP39^*23*^). Thus, Lys170 likely acts as a general base during 2’-O methyl transfer in SARS-CoV-2. One possible role of the water molecule is that it co-operates with Asp130 to transiently stabilize the transfer of the positively charged methyl group from SAM to 2’-O of A_1_ base. We propose that once occupied by SAM, the catalytic pocket is primed for A_1_ base binding and catalysis. This may be a general mechanism for 2’-O methylation for all coronaviruses. Although all components required for methylation are present in our crystals, we have not observed methylation of 2’-OH, possibly due to a requirement for additional RNA sequence downstream of the target A_1_ base^*8*^. Thus, the ternary structure that we have described may represent a pre-reaction state of 2’-O methyl transfer.

### Acquired mutations in SARS-CoV-2 and their implications

Next, to better understand the role of acquired mutations in SARS-CoV-2 nsp16 (compared to CoV-1), we aligned nsp16 sequences from 9 coronavirus strains representing all three sub classes (α, β, and *γ*), and mapped their positions in the current structure (Supplementary Fig. 1). Although most residues that participate in catalysis and substrate/SAM binding are conserved across species, notable differences were found (Supplementary Fig. 1, 3B). Compared to CoV-1, acquired mutations span the entire nsp16 sequence: E32D and N33S (gate loop1); K135R (gate loop 2); and S188A, A209C, T223V, N265G, Y272L, E276S, V290I, and I294V (Supplementary Fig. 1, 3B). Given the role of gate loop regions in RNA substrate binding, mutations E32D and N35S in gate loop 1 and K135R in gate loop 2 may influence RNA binding or enzyme kinetics.

Three specific mutations – S188A (β8), A209C (β9), and E276S (α6) forms part of the adenosine binding pocket. Two of the mutations A209C (β9), and E276S (α6) are not present in any other CoV strain, whereas S188A, is present in one human strain (HKU1), two avian strains (TCoV and IBV) and one mouse strain (MHV) (Fig. 3, Supplementary Fig. 3). At ~ 25 Å away, and on the back of the catalytic center, this ligand binding pocket does not present obvious characteristics for adenine specificity such as an aspartic acid or asparagine residue, which would usually make a hydrogen bond with the N6 amino group of adenines. Nonetheless, it displays two similarities with the SAM binding pocket: the purine ring in both pockets resides in an acidic environment, and both pockets harbor a cysteine near N6 of the purine ring. In the SAM pocket, Cys115 is 4.1 Å from the N6 amino group and positioned at a ~ 64º angle between N1 and the C6 atoms of the adenine ring with respect to Cys115 (Fig. 2C). An equivalent cysteine in the adenosine pocket, Cys209, resides at 3.5 Å, and at a ~ 85º angle from the N6 of adenine. The purine ring of adenosine is further stabilized by stacking interactions with Trp198 and Thr58, and its 5’ tail, while exposed to solvent, interacts with Asn13 (Fig. 3A). Thus, the lack of extensive interactions for adenosine suggests that this pocket is somewhat non-specific but can accommodate small molecules with a heterocyclic ring. Interestingly, a binary structure of SARS-CoV-2 nsp16 (PDB ID: 6W4H) shows a sugar moiety (β-D-fructopyranose) occupying the position of adenosine. In SARS-CoV-1 nsp16 (PDB ID: 2XYR)^*14*^, a metal ion (Mg^2+^) is present in vicinity (~4 Å) of the adenosine or β-D-fructopyranose ligands binding site (**Supplementary Fig. 4**). Thus, we propose that this distant region endows nsp16 with the unique ability to bind small molecules outside of the catalytic pocket. Future studies will clarify the impact of this binding region on nsp16 function and viral pathogenesis.

### Concluding Remarks

In conclusion, we have presented here a high-resolution structure of a ternary complex of nsp16/nsp10 in the presence of an RNA cap and SAM, captured just prior to ribose 2’-O methylation. The structure reveals key conformational changes in the RNA substrate binding site, and additional biochemical determinants for the catalysis of RNA Cap methylation. Our structure also reveals a potential allosteric site which may be targeted in the development of antiviral therapies to treat SARS-CoV-2 infections.

## METHODS SUMMARY

The nsp16/nsp10 enzyme complex was expressed and purified from *E. coli*. The complex was crystallized in the presence of an RNA cap analogue and S-adenosyl methionine (SAM) and adenosine. The crystals belong to the space group P3_1_21 with unit cell dimensions of a=b=168 Å, c=52.3Å, and α=β=90°, γ=120. They diffract X-rays to 1.8 Å resolution with synchrotron radiation. The structure was solved with molecular replacement using a binary complex (nsp16/nsp10/SAM; PDB ID 3R24) as a template. The final model was refined to 1.8 Å resolution with R_free_ and R_work_ values of ~ 19.1% and 17.2%, respectively (Supplementary Table 1).

## Acknowledgments

We are grateful to beamline scientists at NECAT-24ID, APS, Chicago for providing synchrotron beamtime and facilitating data collection. We thank Drs. Bruce Nicholson and Patrick Sung and for critically reading the manuscript. This work was supported by funding from the Max and Minnie Tomerlin Voelcker Foundation, San Antonio Area Foundation, a Rising STARs award from the UT System, a Pilot award from the UT Health San Antonio (UTHSA), and laboratory startup funds from the Greehey Children’s Cancer Research Institute (GCCRI) of UTHSA. S.Q. is supported by the GCCRI. T.V. is supported by a Research Training Award (RP170345) from the Cancer Prevention Research Institute of Texas (CPRIT). R.A.H. is supported by NIH CA205224. We also thank X-ray crystallography core of the UTHSA. This work is based on research conducted at the Northeastern Collaborative Access Team beamlines, which are funded by NIH grant P30GM124165, and U.S. Department of Energy (DOE) Contract DE-AC02-06CH11357.

## Author Contributions

Y.K.G. conceived, designed, supervised the study, and performed crystallographic studies; S.A. and T.V. expressed and purified proteins; T.V. performed crystallographic studies; S.Q. and T.V. performed biochemical assays; S.-H.C and N.D. performed enzymatic assays; R.A.H., L.M.-S., J.P., and F.O. provided reagents. Y.K.G. and S.-H.C. wrote the manuscript.

## Author Information

Files for atomic coordinates and structure factors were deposited in the Protein Data Bank under accession codes 6WKS. Correspondence and requests for material should be addressed to Y.K.G.

## METHODS

### Protein expression and purification

The coding sequences of nsp16 (NCBI reference sequence YP_009725311.1) and nsp10 (NCBI reference sequence: YP_0009725306.1) of the Wuhan seafood market pneumonia virus isolate Wuhan-Hu-1 (NC_045512) were cloned into a single pETduet-1 vector downstream to a 6XHis tag sequence. This plasmid was transformed into an *E. coli* expression strain BL21 (DE3). The transformed cells were grown in Terrific Broth medium supplemented with ampicillin at 37 °C. Protein expression was induced by adding 0.4 mM isopropyl β-D-thiogalactopyranoside (IPTG) at OD_600_ = 0.6 followed by continued incubation of the cultures for 14 hours at 18°C. Cells were then harvested by centrifugation at 6000 rpm for 20 minutes. Thereafter, cells were re-suspended in ice-cold lysis buffer. Cell lysis was accomplished using a microfluidizer (Analytik, UK). The lysate was centrifuged at 40,000 rpm for 40 minutes and passed through a 0.22µm filter. The clarified soluble fraction was loaded on to a Nuvia IMAC column (Bio Rad) pre-equilibrated in binding buffer containing 25 mM Tris-HCl pH 8.0, 0.5 M NaCl, 0.1mM TCEP, 10% Glycerol. The proteins were eluted by increasing the concentration of imidazole. The 6XHis tag was then proteolytically removed.

The tag-free sample was re-applied to the IMAC column to separate the un-cleaved protein fraction. The proteins were purified by successive passage through HiTrap heparin and Superdex 75 columns (GE Healthcare). The nsp16/nsp10 complex was eluted as a single homogenous species in a final buffer containing 25 mM Tris-HCl pH 8.0, 0.5 M NaCl, 0.1mM TCEP, 10% glycerol and 5 mM MgSO_4_. The purified protein complex was concentrated to 5 mg/mL, mixed with 5 molar excess of adenosine, and subjected to extensive crystallization trials.

### Crystallization

Initial crystals were grown by the sitting drop vapor diffusion method. After 3-4 rounds of optimizations by varying pH, precipitant and salt concentrations, we grew larger crystals amenable to synchrotron radiation. We soaked these crystals with ^me7^GpppA RNA analogue representative of Cap-0 structure. Diffraction-quality crystals were grown in a crystallization solution containing 10% (v/v) MPD, 0.1 M HEPES pH 7.0. Before data collection, the co-crystals were cryo-protected by serial soaks in solution containing original mother liquor and increasing concentrations (0 → 20% v/v) of ethylene glycol, and then flash-frozen in liquid nitrogen.

### X-ray diffraction data collection and structure determination

Crystals of the nsp16/nsp10/SAM/Cap-O/adenosine complex diffracted X-rays to 1.8 Å and belong to space group P3_1_21 with unit cell dimensions a=b=168Å, c = 52.3Å, c = 313.9Å, α = β = 90°, and γ = 120°, and with one nsp16/nsp10 heterodimer per asymmetric unit. All data were indexed, integrated, and scaled using XDS^*24*^, aimless^*25*^, and various ccp4 suite programs (truncate, freeflag, and mtz2various)^*26*^. The structure was solved by molecular replacement using the SAM-bound nsp16/nsp10 (PDB ID: 3R24)^*8*^ structure as a template in Phaser ^*26*^. The resulting maps indicated the electron densities for SAM, Cap-0 and adenosine, and regions in nsp16 with extreme conformational changes (gate loops 1 and 2). Ligand restraints were generated using Grade (Global Phasing Ltd.). We iteratively rebuilt and refined the model using the programs Coot^*27*^ and REFMAC^*26*^. The final model was refined to R_factor_ to ~ 17.1% and the R_free_ to ~19%. A Ramachandran plot for the final model shows 93.8% of the residues in the most preferred regions, 4.4% in the additionally allowed regions, and 1.7% in the disallowed regions. All figures that depict structural models were generated using Pymol.

### Determining affinities for nsp16/nsp10 ligands

The purified nsp16/nsp10 protein complex was labeled with a protein labeling Kit (Monolith, RED-NHS 2nd Generation kit, Cat# MO-L011). In brief, 20 µM of protein was incubated with dye solution (60 µM) in the labeling buffer and the reaction was allowed to proceed at room temperature for 30 min. Each ligand was dissolved in the Microscale thermophoresis (MST) reaction buffer containing 20 mM HEPES pH 7.5, 150 mM NaCl, 0.5% glycerol, and 0.05% Tween 20. Two-fold serial dilutions started from 2 mM ligand concentrations were made in 12 steps. The labeled protein at a final concentration of 20 nM was equally mixed into each ligand reaction (ligand concentration range 500 nM to 1 mM). The final reaction mixtures were loaded into premium capillary chips (Monolith Cat# MO-AK005) and measured on a Monolith NT.115 instrument (NanoTemper Technologies) at 20% excitation power and 40% MST power at 25 °C. Results shown here are from two independent experiments. Data were fitted by a single-site binding model in GraphPad Prism (GraphPad Software, San Diego, CA).

### Enzyme activity assay

To assess the relative activity of nsp16/nsp10 on RNA with an adenosine or guanosine as the initiating nucleotide, increasing concentrations of purified nsp16/nsp10 were allowed to react with 1 µM ^me7^Gppp**A**- or ^me7^Gppp**G**-capped 25 nt RNA in a buffer containing 50 mM Tris-HCl, pH 8.0, 5 mM KCl, 1 mM DTT, 0.2 mM SAM and 1 mM MgCl_2_. The reactions were incubated at 37°C for 30 min and stopped by heating at 75°C for 5 min. The reactions were subjected to LC/MS intact mass analysis. Briefly, nucleic acids in the samples were separated using a Thermo DNAPac™ RP Column (2.1 × 50 mm, 4 µm) on a Vanquish Horizon UHPLC System, followed by mass determination using a Thermo Q-Exactive Plus mass spectrometer. The raw data was deconvoluted using Promass (Novatia Inc.). The ratio of the deconvoluted mass peak intensity of the reactants and the expected products were used to estimate the percentage of 2’-O methylation.

The two RNA substrates used in this assay only differ by 1 base (**A** vs **G**) at N_1_ base position: ^me7^Gppp**A**UAGAACUUCGUCGAGUACGCUCAA-[6-FAM] ^me7^Gppp**G**UAGAACUUCGUCGAGUACGCUCAA-[6-FAM]

## SUPPLEMENTARY DATA

**Fig. S1.**
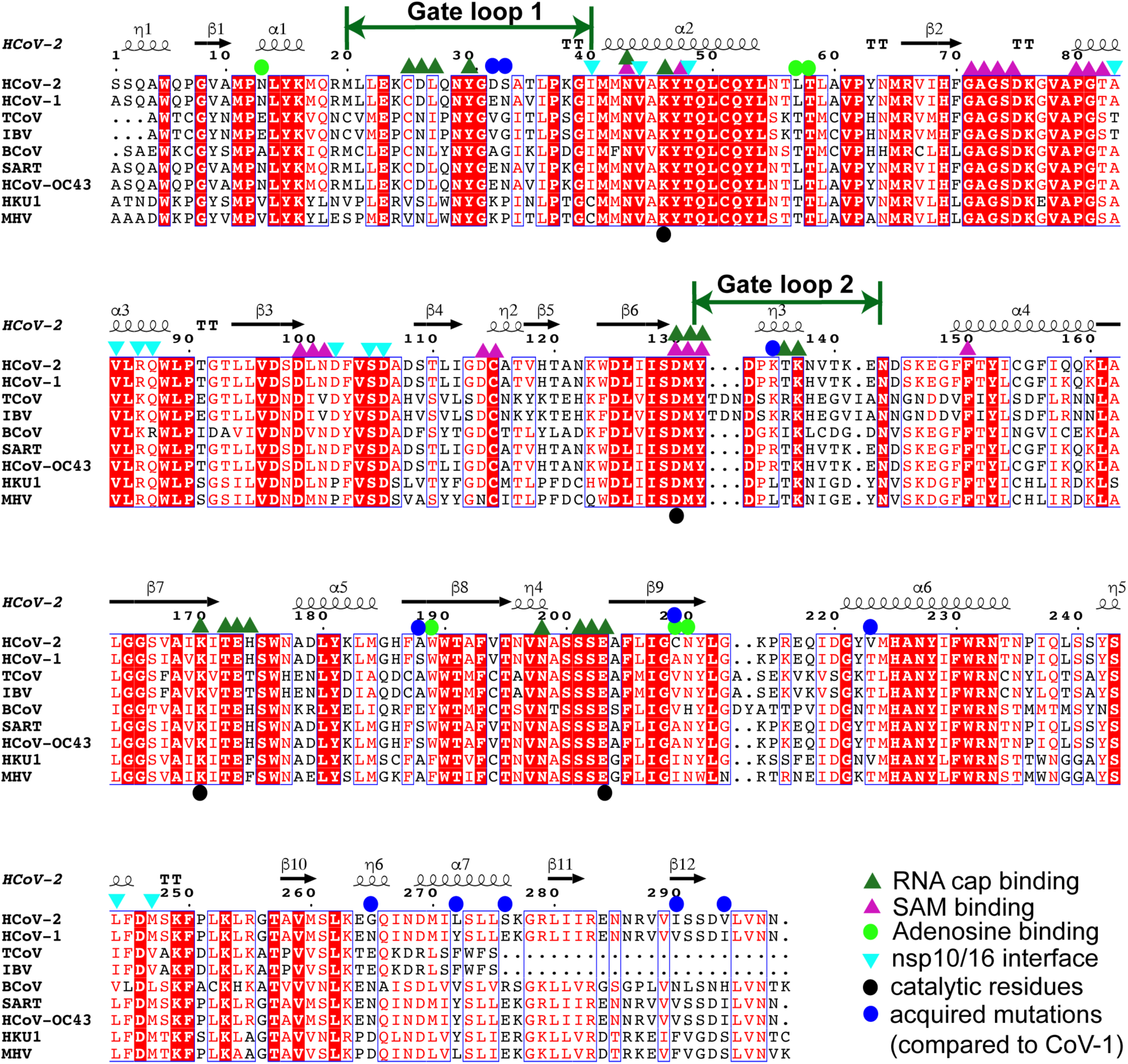
Structural elements of nsp16. The SARS-CoV-2 nsp16 adopts a SAM-MT fold^*17*^. The secondary structural elements are shown above the aligned protein sequences of nsp16 from nine coronavirus members (HCoV-2, SARS-CoV-2, YP_009725311.1; HCoV-1, SARS-CoV-1, Uniprot ID: P0C6×7; TCoV, turkey coronavirus, YP_001941189.1; IBV, infectious bronchitis virus; BCoV, bat coronavirus, YP_008439226.1; SART, human SARS CoV, NP_828873.2; HCoV-OC43, human coronavirus OC43; HKU1, human coronavirus, YP_460023.1; and MHV, murine hepatitis virus, YP_209243.1). The protein sequences were first aligned using MUSCLE server (https://www.ebi.ac.uk/Tools/msa/muscle/). The ESPript server^*25*^ was used to assign secondary structural elements of the SARS-CoV-2 nsp16 structure. The substrate (RNA cap) binding regions span two-thirds of the protein, and the regions belonging to the first half participate in SAM binding. The major contributions to cap binding come from gate loop regions 1 and 2. Different symbols atop residues represent their respective involvement in substrate (green triangles), co-factor (magenta triangles), adenosine (bright green circles), and inter-subunit interactions (inverted triangles in cyan). Catalytic residues are annotated by black circles on the bottom. Positions of acquired mutations in SARS-CoV-2 (compared to SARS-CoV-1) are denoted by blue circles.

**Fig. S2.**
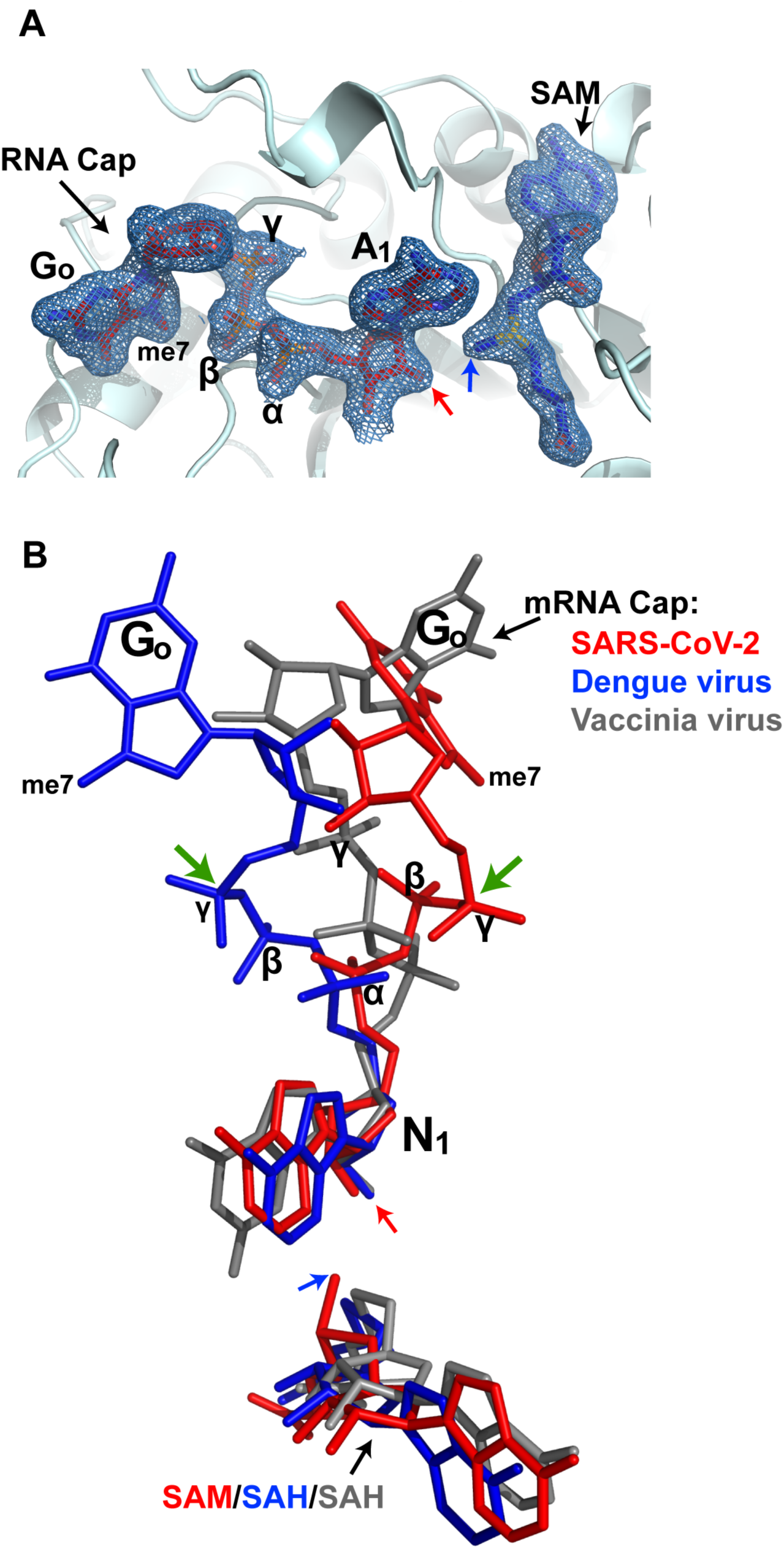
Electron density maps and orientation of RNA cap in different viruses. A close-up view of the binding pockets for substrate (RNA cap, red stick) and cofactor (SAM, blue stick) analogues. The 2Fo-Fc electron density map for these ligands contoured at 1σ is shown as a blue mesh. Positions of the methyl group (in donor SAM), and the acceptor moiety (2’-O in ribose of the target nucleotide A_1_) are depicted by blue and red arrows, respectively. B) A superposition of N_1_ ribose of RNA caps of SARS-CoV-2 (present work, red), Dengue virus (PDB ID: 5DTO, blue), and vaccinia virus (PDB ID: 1AV6, grey) shows 180º rotation in γ-phosphates (green arrows) of RNA caps of SARS-CoV-2 and Dengue viruses.

**Figure S3.**
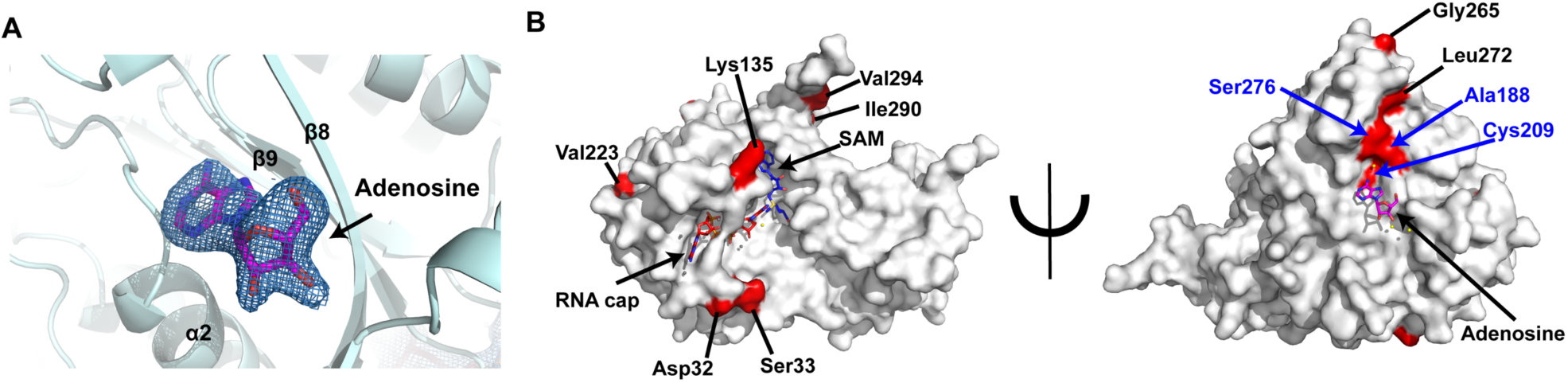
Adenosine binding and mapping of acquired mutations. A close-up view of the binding pockets for adenosine (magenta stick) analogue. The 2Fo-Fc electron density map contoured at 1σ is shown around the adenosine as a blue mesh. B) Positions of acquired mutations in SARS-CoV-2 (red surface) compared to from SARS-CoV-1 are mapped on nsp16 surface (light grey). The allosteric triad (Ala88, Cys209, Ser276) residues specific to SARS-CoV-2 nsp16 are labeled and marked in blue.

**Figure S4.**
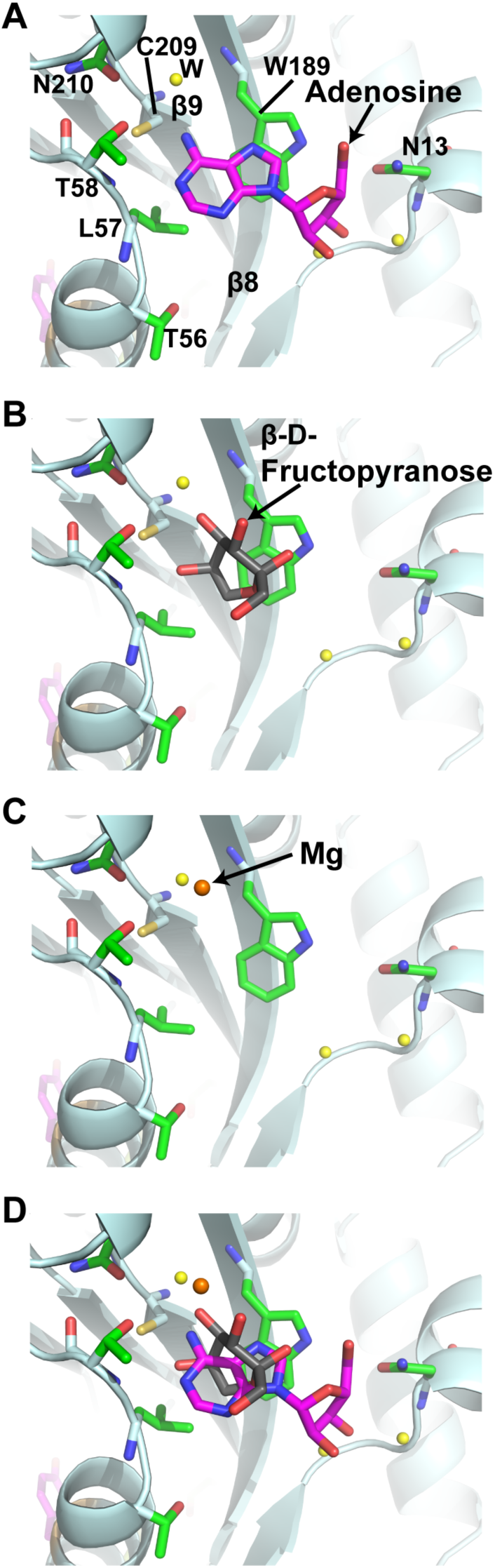
Putative allosteric site in SARS-CoV-2 nsp16. A) Binding mode of adenosine (magenta) in ternary structure of SARS-CoV-2 nsp16. B) Binding mode of β-D-fructopyranose (grey) in binary structure of SARS-CoV-2 nsp16 (PDB ID: 6W4H). C) Binding of Mg^2+^ (orange sphere) in binary structure of SARS-CoV-1 nsp16 (PDB ID: 2XYR). D) An overlay of all 3 ligands in the putative allosteric site. The protein structure of only SARS-CoV-2 nsp16 is shown in all panels.

**Supplementary Table 1.** Data Collection and Refinement Statistics

|  |  |
| --- | --- |
|  | nsp16/nsp10/SAM/<sup>me7</sup>G<sub>o</sub>pppA |
| **Data Collection** |  |
| Space Group | P3<sub>1</sub>21 |
| Cell dimensions |  |
| a=b, c (Å) | 168, 52.3 |
| α=β, γ (°) | 90, 120 |
| Wavelength(Å) | 0.9792 |
| Resolution (Å) | 1.8 |
| R<sub>sym</sub> (%)<sup>a,b</sup> | 13.71 (165) |
| Number of reflections | 1677960 (74259) |
| Completeness | 99.9 (100) |
| Redundancy | 20.8 (20.2) |
| I/sigma | 14.15 (1.35) |
| **Refinement** |  |
| Resolution (Å) | 30-1.8 |
| R<sub>factor</sub> (%)<sup>c</sup> / R<sub>free</sub> (%)<sup>d</sup> | 17.2 / 19.1 |
| *Nonhydrogen atoms* |  |
| Protein / ligands / Water | 3526 / 114 / 252 |
| *Average B-factors (Å<sup>2</sup>)* |  |
| Protein / ligands / Water | 42.9 / 43.1 / 47.8 |
| *R.m.s. deviations* |  |
| Bond lengths (Å) | 0.015 |
| Bond angles (°) | 1.89 |
| *Ramachandran plot* |  |
| Most Favored (%) | 93.8 |
| Additional allowed (%) | 4.6 |
| Disallowed (%) | 1.4 |

<sup>a</sup>Values for outermost shells are given in parentheses.
<sup>b</sup>R<sub>sym</sub> = $\sum |I - \langle I \rangle| / \sum I$ , where I is the integrated intensity of a given intensity.
<sup>c</sup>R<sub>factor</sub> = $\sum ||F_{\text{observed}}| - |F_{\text{calculated}}|| / \sum |F_{\text{observed}}|$ .
<sup>d</sup>R<sub>free</sub> was calculated using 5.3% of random data omitted from the refinement

## REFERENCES

[1] (2020) COVID-19 Coronavirus Pandemic, In https://www.worldometers.info/coronavirus/.

[2] Zhou, P., Yang, X. L., Wang, X. G., Hu, B., Zhang, L., Zhang, W., Si, H. R., Zhu, Y., Li, B., Huang, C. L., Chen, H. D., Chen, J., Luo, Y., Guo, H., Jiang, R. D., Liu, M. Q., Chen, Y., Shen, X. R., Wang, X., Zheng, X. S., Zhao, K., Chen, Q. J., Deng, F., Liu, L. L., Yan, B., Zhan, F. X., Wang, Y. Y., Xiao, G. F., and Shi, Z. L. (2020) A pneumonia outbreak associated with a new coronavirus of probable bat origin, Nature 579, 270–273.

[3] Cencic, R., Desforges, M., Hall, D. R., Kozakov, D., Du, Y., Min, J., Dingledine, R., Fu, H., Vajda, S., Talbot, P. J., and Pelletier, J. (2011) Blocking eIF4E-eIF4G interaction as a strategy to impair coronavirus replication, J Virol 85, 6381–6389.

[4] Bouvet, M., Debarnot, C., Imbert, I., Selisko, B., Snijder, E. J., Canard, B., and Decroly, E. (2010) In vitro reconstitution of SARS-coronavirus mRNA cap methylation, PLoS Pathog 6, e1000863.

[5] Bouvet, M., Imbert, I., Subissi, L., Gluais, L., Canard, B., and Decroly, E. (2012) RNA 3’-end mismatch excision by the severe acute respiratory syndrome coronavirus nonstructural protein nsp10/nsp14 exoribonuclease complex, Proc Natl Acad Sci U S A 109, 9372–9377.

[6] Decroly, E., Ferron, F., Lescar, J., and Canard, B. (2011) Conventional and unconventional mechanisms for capping viral mRNA, Nat Rev Microbiol 10, 51–65.

[7] Ivanov, K. A., and Ziebuhr, J. (2004) Human coronavirus 229E nonstructural protein 13: characterization of duplex-unwinding, nucleoside triphosphatase, and RNA 5’-triphosphatase activities, J Virol 78, 7833–7838.

[8] Chen, Y., Su, C., Ke, M., Jin, X., Xu, L., Zhang, Z., Wu, A., Sun, Y., Yang, Z., Tien, P., Ahola, T., Liang, Y., Liu, X., and Guo, D. (2011) Biochemical and structural insights into the mechanisms of SARS coronavirus RNA ribose 2’-O-methylation by nsp16/nsp10 protein complex, PLoS Pathog 7, e1002294.

[9] Zust, R., Cervantes-Barragan, L., Habjan, M., Maier, R., Neuman, B. W., Ziebuhr, J., Szretter, K. J., Baker, S. C., Barchet, W., Diamond, M. S., Siddell, S. G., Ludewig, B., and Thiel, V. (2011) Ribose 2’-O-methylation provides a molecular signature for the distinction of self and non-self mRNA dependent on the RNA sensor Mda5, Nat Immunol 12, 137–143.

[10] Daffis, S., Szretter, K. J., Schriewer, J., Li, J., Youn, S., Errett, J., Lin, T. Y., Schneller, S., Zust, R., Dong, H., Thiel, V., Sen, G. C., Fensterl, V., Klimstra, W. B., Pierson, T. C., Buller, R. M., Gale, M., Jr., Shi, P. Y., and Diamond, M. S. (2010) 2’-O methylation of the viral mRNA cap evades host restriction by IFIT family members, Nature 468, 452–456.

[11] Hyde, J. L., and Diamond, M. S. (2015) Innate immune restriction and antagonism of viral RNA lacking 2-O methylation, Virology 479-480, 66–74.

[12] Menachery, V. D., Yount, B. L., Jr., Josset, L., Gralinski, L. E., Scobey, T., Agnihothram, S., Katze, M. G., and Baric, R. S. (2014) Attenuation and restoration of severe acute respiratory syndrome coronavirus mutant lacking 2’-o-methyltransferase activity, J Virol 88, 4251–4264.

[13] Wang, Y., Sun, Y., Wu, A., Xu, S., Pan, R., Zeng, C., Jin, X., Ge, X., Shi, Z., Ahola, T., Chen, Y., and Guo, D. (2015) Coronavirus nsp10/nsp16 Methyltransferase Can Be Targeted by nsp10-Derived Peptide In Vitro and In Vivo To Reduce Replication and Pathogenesis, J Virol 89, 8416–8427.

[14] Decroly, E., Debarnot, C., Ferron, F., Bouvet, M., Coutard, B., Imbert, I., Gluais, L., Papageorgiou, N., Sharff, A., Bricogne, G., Ortiz-Lombardia, M., Lescar, J., and Canard, B. (2011) Crystal structure and functional analysis of the SARS-coronavirus RNA cap 2’-O-methyltransferase nsp10/nsp16 complex, PLoS Pathog 7, e1002059.

[15] de Clercq, E., and Montgomery, J. A. (1983) Broad-spectrum antiviral activity of the carbocyclic analog of 3-deazaadenosine, Antiviral Res 3, 17–24.

[16] Montgomery, J. A., Clayton, S. J., Thomas, H. J., Shannon, W. M., Arnett, G., Bodner, A. J., Kion, I. K., Cantoni, G. L., and Chiang, P. K. (1982) Carbocyclic analogue of 3-deazaadenosine: a novel antiviral agent using S-adenosylhomocysteine hydrolase as a pharmacological target, J Med Chem 25, 626–629.

[17] Votruba, I., Holy, A., and De Clercq, E. (1983) Metabolism of the broad-spectrum antiviral agent, 9-(S)-(2,3-dihydroxypropyl) adenine, in different cell lines, Acta Virol 27, 273–276.

[18] Coloma, J., Jain, R., Rajashankar, K. R., Garcia-Sastre, A., and Aggarwal, A. K. (2016) Structures of NS5 Methyltransferase from Zika Virus, Cell Rep 16, 3097–3102.

[19] Hodel, A. E., Gershon, P. D., Shi, X., and Quiocho, F. A. (1996) The 1.85 A structure of vaccinia protein VP39: a bifunctional enzyme that participates in the modification of both mRNA ends, Cell 85, 247–256.

[20] Martin, J. L., and McMillan, F. M. (2002) SAM (dependent) I AM: the S-adenosylmethionine-dependent methyltransferase fold, Curr Opin Struct Biol 12, 783–793.

[21] Joseph, J. S., Saikatendu, K. S., Subramanian, V., Neuman, B. W., Brooun, A., Griffith, M., Moy, K., Yadav, M. K., Velasquez, J., Buchmeier, M. J., Stevens, R. C., and Kuhn, P. (2006) Crystal structure of nonstructural protein 10 from the severe acute respiratory syndrome coronavirus reveals a novel fold with two zinc-binding motifs, J Virol 80, 7894–7901.

[22] Zhao, Y., Soh, T. S., Lim, S. P., Chung, K. Y., Swaminathan, K., Vasudevan, S. G., Shi, P. Y., Lescar, J., and Luo, D. (2015) Molecular basis for specific viral RNA recognition and 2’-O-ribose methylation by the dengue virus nonstructural protein 5 (NS5), Proc Natl Acad Sci U S A 112, 14834–14839.

[23] Hodel, A. E., Gershon, P. D., and Quiocho, F. A. (1998) Structural basis for sequence-nonspecific recognition of 5’-capped mRNA by a cap-modifying enzyme, Mol Cell 1, 443–447.

[24] Kabsch, W. (2010) Xds, Acta Crystallogr D Biol Crystallogr 66, 125–132.

[25] Evans, P. R. (2007) An introduction to stereochemical restraints, Acta Crystallogr D Biol Crystallogr 63, 58–61.

[26] Collaborative Computational Project, N. (1994) The CCP4 suite: programs for protein crystallography, Acta Crystallogr D Biol Crystallogr 50, 760–763.

[27] Emsley, P., and Cowtan, K. (2004) Coot: model-building tools for molecular graphics, Acta Crystallogr D Biol Crystallogr 60, 2126–2132.

## References

[1] Martin, J. L., and McMillan, F. M. (2002) SAM (dependent) I AM: the S-adenosylmethionine-dependent methyltransferase fold, Curr Opin Struct Biol 12, 783–793.

[2] Robert, X., and Gouet, P. (2014) Deciphering key features in protein structures with the new ENDscript server, Nucleic Acids Res 42, W320–324.

